# Metabolomic-based biomarker discovery for non-invasive lung cancer screening: A case study

**DOI:** 10.1101/063321

**Authors:** Keiron O’Shea, Simon J.S. Cameron, Keir E Lewis, Chuan Lu, Luis AJ Mur

**Author notes:** Corresponding author *Email address:* (Luis AJ Mur).

## Abstract

**Background:** Lung cancer (LC) is one of the leading lethal cancers worldwide, with an estimated 18.4% of all cancer deaths being attributed to the disease. Despite developments in cancer diagnosis and treatment over the previous thirty years, LC has seen little to no improvement in the overall five year survival rate after initial diagnosis.

**Methods:** In this paper, we extended a recent study which profiled the metabolites in sputum from patients with lung cancer and age-matched volunteers smoking controls using flow infusion electrospray ion mass spectrometry. We selected key metabolites for distinguishing between different classes of lung cancer, and employed artificial neural networks and leave-one-out cross-validation to evaluate the predictive power of the identified biomarkers.

**Results:** The neural network model showed excellent performance in classification between lung cancer and control groups with the area under the receiver operating characteristic curve of 0.99. The sensitivity and specificity of for detecting cancer from controls were 96% and 94% respectively. Furthermore, we have identified six putative metabolites that were able todiscriminate between sputum samples derived from patients suffering small cell lung cancer (SCLC) and non-small cell lung cancer. These metabolites achieved excellent cross validation performance with a sensitivity of 80% and specificity of 100% for predicting SCLC.

**Conclusions:** These results indicate that sputum metabolic profiling may have potential for screening of lung cancer and lung cancer recurrence, and may greatly improve effectiveness of clinical intervention.

## 1. Introduction

The year 2008 saw an estimated 12.7 million new cases of cancer, and 7.6 million cancer-related deaths worldwide [1]. While the incidence and mortality rates of most cancers is decreasing in the developed world, they are rising in emerging economies such as China and India. Migrant studies have found that cancer rates in the descendent generation of migrants tends to shift toward the host country, suggesting that environmental risk factors such as smoking and weight are responsible for the global variance in cancer rates [2].

### 1.1 Lung cancer

Lung cancer is a major cause of death in the developed and developing worlds. It is the leading cause of cancer-related deaths in men, and second only to breast cancer in women. There was an estimated 1.6 million new cases of lung cancer and 1.4 million deaths in 2008. This accounts for 12.6% of all cancer incidence and a staggering 18.4% of all cancer-related deaths [2]. This can be attributed to its poor prognosis, with the five-year survival rate being a mere 15%. Despite recent advances in lung cancer treatment, survival rates are low when compared to other forms of cancer [3]. However, improvements in surgical techniques and chemotherapy over the past twenty years has resulted in one-year lung cancer survival rates drastically improving. Despite this, the overall five-year lung cancer survival rates have remained stagnant at 6% for small cell lung cancer and 18% for non-small cell lung cancer. Unfortunately the vast majority (85%) of lung cancer cases are diagnosed at advanced stages, heavily reducing the effectiveness of treatment [1]

This can be attributed to the difficulty of effectively diagnosing cancer of the lung at stage early enough to make a real impact. One of the main difficulties is that symptoms of the conditions are often identical to less serious conditions. This makes the pre-clinical diagnosis of lung cancer particularly problematic as the observed symptoms are often confused with other respiratory conditions. Prognostic factors may help diagnose patients who show symptoms of a disease, or have an increased chance of recurrence or progression to advanced disease which should support clinicians in the creation of appropriate treatment plans. The World Health Organisation (WHO) have set out ten key principles to be met by an effective screening procedure in order for it to be beneficial and cost effective [4]. Currently there are no lung cancer screening techniques of which meet all of the ten conditions laid out by the WHO.

### 1.2 Metabolomic insights into lung cancer

An emerging screening methodology to other traditional screening methods is the utilisation of molecular biomarkers in biofluids. The ease of analysis of biofluids using mass spectrometry (MS) or nuclear magnetic resonance (NMR) makes metabolomics a well-suited methodology for the non-invasive detection of biomarkers in lung cancer. Current focus of metabolomics in lung cancer has been on the exploitation of serum, urine and tumour biopsies. For example, the analysis of serum using liquid chromatography (LC-MS) and gas chromatography (GC-MS) approaches have suggested a potential use for biomarkers of lung cancer. A small-scale pilot study sampling lung cancer patients before and after surgical intervention, alongside patients without lung cancer has suggested ten candidate biomarkers for lung cancer, including sphingosine, oleic acid and serine [5].

Sputum has been suggested as a potential biofluid source of biomarkers in lung cancer [3, 6]. Recent work has used Fourier Transform Infra-Red (FTIR) spectroscopy as a non-invasive method to detect lung cancer in sputum samples. This work concluded that FTIR was able to sufficiently distinguish between lung cancer and control samples, and effectively act as a non-invasive, high-throughput and cost-effective method for screening sputum samples from high-risk patients. Furthermore, it further validated the use of sputum as an effective biofluid for lung cancer screening [7].

### 1.3 Artificial Neural Networks

Artificial neural networks (ANNs) are a class of sophisticated computational modelling structures that are inspired by biological neurological systems, regarding how they are able to learn and process highly non-linear information [37]. The past three decades have seen ANNs being widely used for biomedical decision support systems [8, 9, 10, 11, 12].

In general, an artificial neural network is formed of interconnected processing units, commonly referred to as neurons. Each neuron applies an activation function over the weighted sum of the incoming stimuli (or inputs), and generate an output signal, which could be the input signal for other neurons. Many different neural network architectures exist, in this paper we will focus on the popular feed-forward artificial neural network, in particular the multi-layer perceptron (MLP) [13], that usually consists of multiple layers of neurons – the input layer, one or more hidden layers and the final output layer, as illustrated in Figure 1.

**Figure 1:**
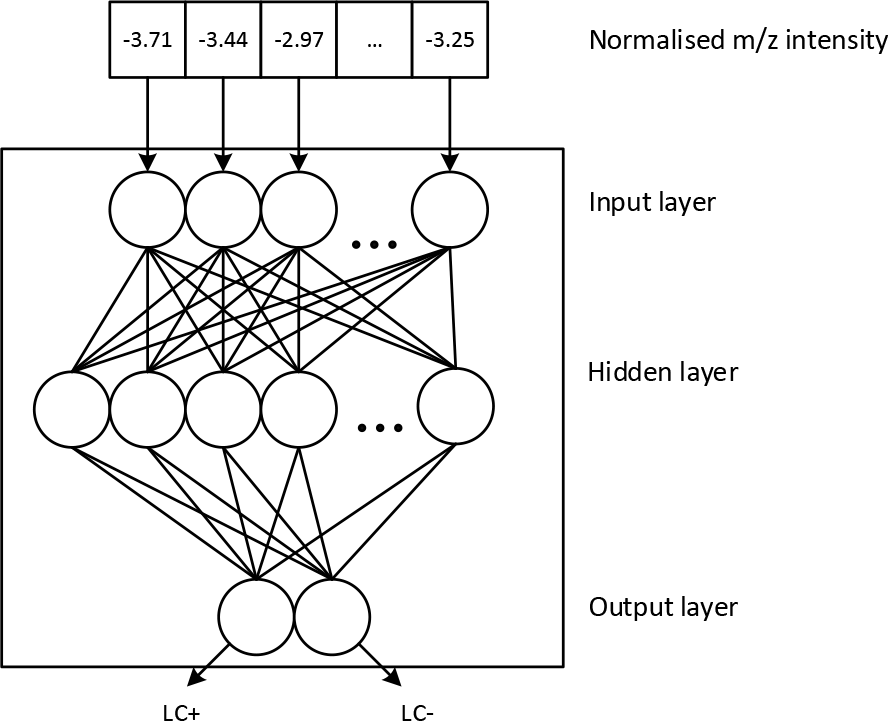
Illustration of a three-layered feed-forward artificial neural network, where each neuron in one layer has connections to the subsequent layer.

The design of network architectures involves setting the number of hidden layers, the number of neurons within each layer, the connections between them, and the type of activation function to use. The connection weights in the network could be adjusted through a learning algorithm that minimises the amount of error in the outputs compared to the true ones. Generalisation, and to avoid overfitting the training data would be a central issue both in network design and training.

## 2. Case Study

We have recently developed this approach by employing flow-infusion electrospray-Mass Spectrometry (FIE-MS) to evaluate the potential of spontaneous sputum as a source of non-invasive metabolomic biomarkers for LC status [14]. Spontaneous sputum was collected and processed from 34 patients suspected of having LC, and 33 healthy controls. Of the 34 patients, 23 were subsequently diagnosed with LC (LC+) at various stages of disease progression. The clinical characteristics of all samples taken are summarised in Table 1.

**Table 1:**
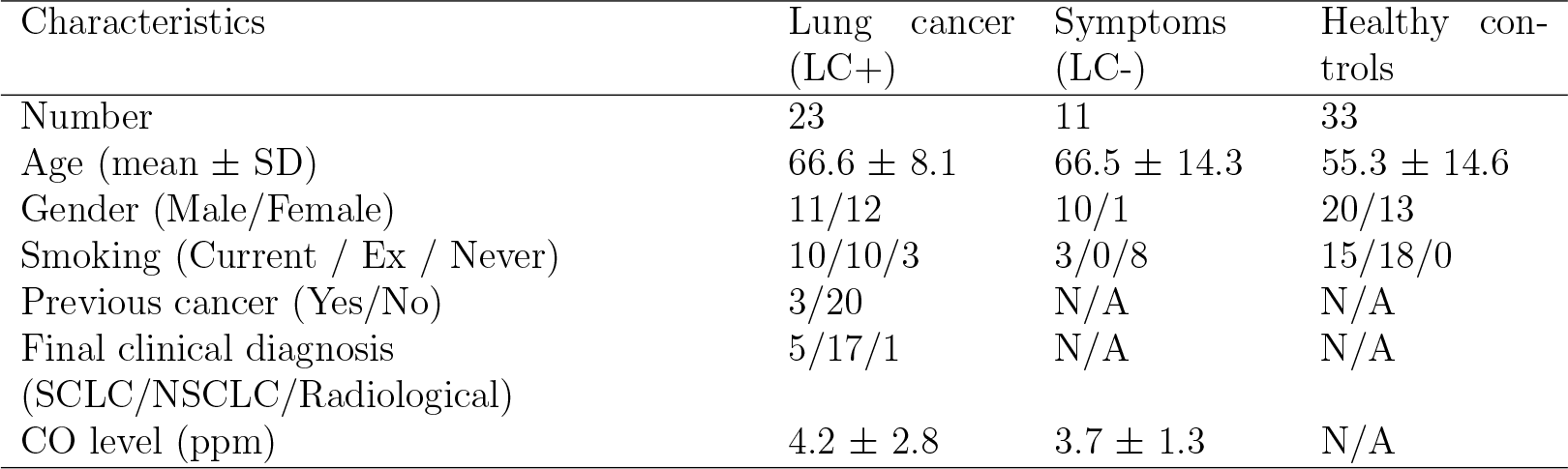
Summary of clinical characteristics of patients with Lung Cancer (LC+), Symptoms (LC−) and Healthy Controls

In these preliminary analyses, discriminatory metabolites were identified using ANOVA and Random Forest and included Ganglioside GM1 which has previously been linked to lung cancer [15]. This suggested that the use of sputum as a non-invasive source of metabolite biomarkers may aid in the development of an at-risk population screening programme for lung cancer or enhanced clinical diagnostic pathways. We now demonstrate how further data-mining of the FIE-MS data has revealed further metabolite biomarkers, and evaluate further the use of metabolomics to yield biomarkers for distinguishing lung cancer type.

### 2.1 Ethics statement

The MedLung observational study (UKCRN ID 4682) received loco-regional ethical approval from the Hywel Dda Health Board (05/WMW01/75). Written informed consent was obtained from all participants at least 24 hours before sampling, at a previous clinical appointment, and all data was link anonymised before analysis.

### 2.2 Mass spectrometry

Frozen sputum samples were thawed before being exposed to 0.5 mL of dithiothreitol (DTT) to isolate sputum cells. Each sample was mixed using a vortex mixer for 15 minutes before being centrifuged at 1800g for 10 minutes before removing the supernatant. Sputum pellets were then analysed using Flow Infusion Electrospray Ion Mass spectrometry (FIE-MS).

Signals identified under 50 *m*/*z* were removed, and the resulting FIE-MS data matrix contains 2,582 *m*/*z* values after binning. The data was further preprocessed by total ion count (TIC) normalisation (to ensure the intensity values for each spectrum sum up to one), followed by log10 transformation prior to further data analysis.

### 2.3 Effect of clinical characteristics on the metabolic profiles

To explore the possible effects of the clinical characteristics on the global metabolic profiles, we conducted the so called 50-50 MANOVA test, which is essentially a variant of classical MANOVA that can handle multiple highly correlated responses [16]. We found no significant effect of age, gender or the CO level on the preprocessed metabolomic data (with *p*-values of 0.4, 0.08, and 0.8, respectively), whilst the effect of disease status (LC+/LC− /Control) is really strong (*p*-value = 1e-12). And perhaps not surprisingly, as tobacco smoking is an important risk factor for lung cancer, there is indeed a significant effect of the smoking pack numbers per year over the metabolic profiles for the patients of LC− and LC+ (*p*-value = 2e-5).

### 2.4 Diagnostic modelling with artificial neural networks

To find discriminatory *m*/*z* features, Welch's unequal variance *t*-test have been performed using the pre-processed intensity values after log-transformation. Random forests have also been tried (results not shown), and the top ranked features identified for both methods are quite similar.

Then an ANN was used as a diagnostic model for various binary classification problems, taking the selected *m*/*z* signals (the preprocessed intensity values after log-transform) as the inputs and estimating the probability for individual classes. The activation function was set to hyperbolic tangent for both hidden and output layer; and the number of hidden layers was set to two for all problems based on our initial analysis. Regularisation techniques such as weight decay [17] have been employed to control the complexity of the model parameters in order to avoid overfitting the models to the training data.

The predictive power of neural network classifiers was evaluated using Receiver Operating Characteristic (ROC) analysis and through leave-one out (LOO) cross-validation (CV) - the overall ROC curve and the area under the curve (AUC) from CV was obtained using the pooled test examples from CV. For each binary classification problem and within each round of training, a *t*-test would be performed on the training set, only the *m*/*z* signals with resulting *p*-values < 0.05 were selected as input features for ANN modelling.

## 3. Results and Discussion

Representative spectra of the samples from the LC+, LC− and healthy control sample groups are shown in Figure 2. FIE-MS profiles were analysed using principal component analysis (PCA) (Figure 4). One can observe no clear separation between clinically collected sample groups (LC+ and LC−) if all m/z signals were used for PCA.

Welch *t*-tests provided 445 distinctive *m*/*z* values for LC+ versus healthy controls, and 90 significant *m*/*z* values for LC+ versus LC− from our pooled leave one out cross validation t-tests. PCA of both stratifications showed good discriminative ability when using the pooled features, as shown in Figure 4. The number of neurons in each hidden layer was chosen through grid search [18]. The best-performing models and their diagnostic performance from LOOCV can be found in Table 2. And Figure 6 shows the resulting ROC curves.

**Table 2:** Results of cross-validation prediction performance of our ANN models. The diagnostic performance (except for AUC) was obtained by using the default probability cutoff value of 0.5 to determine the class labels.

| Classification | Mean no. of inputs | No. of hidden neurons | Sensitivity | Specificity | Positive Predictive Value | Negative Predictive Value | AUC |
| --- | --- | --- | --- | --- | --- | --- | --- |
| LC+ (vs Control) | 1730.2 | 100, 50 | 96% | 94% | 92% | 97% | 0.99 |
| LC+ (vs LC-) | 71.9 | 40, 10 | 100% | 91 % | 96% | 100% | 1.00 |
| SCLC (vs NSCLC) | 77.8 | 50, 20 | 80% | 100% | 100% | 94% | 1.00 |

**Figure 2:**
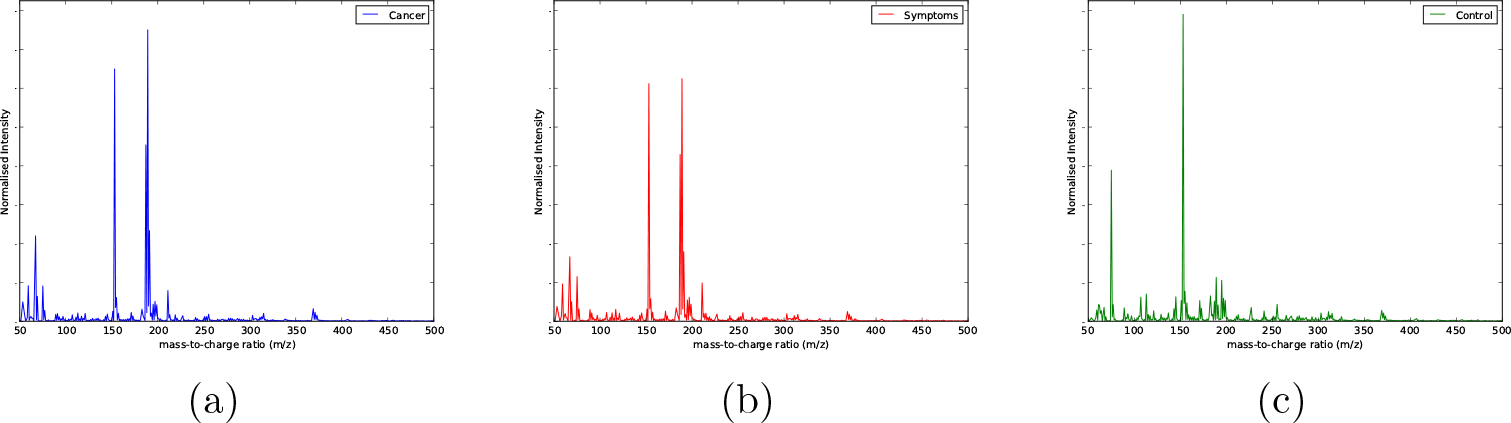
Typical FIE-MS spectra of sputum obtained from sputum obtained from (a) a patient with lung cancer, (b) a patient with symptoms of lung cancer and (c) a healthy control sample.

**Figure 3:**
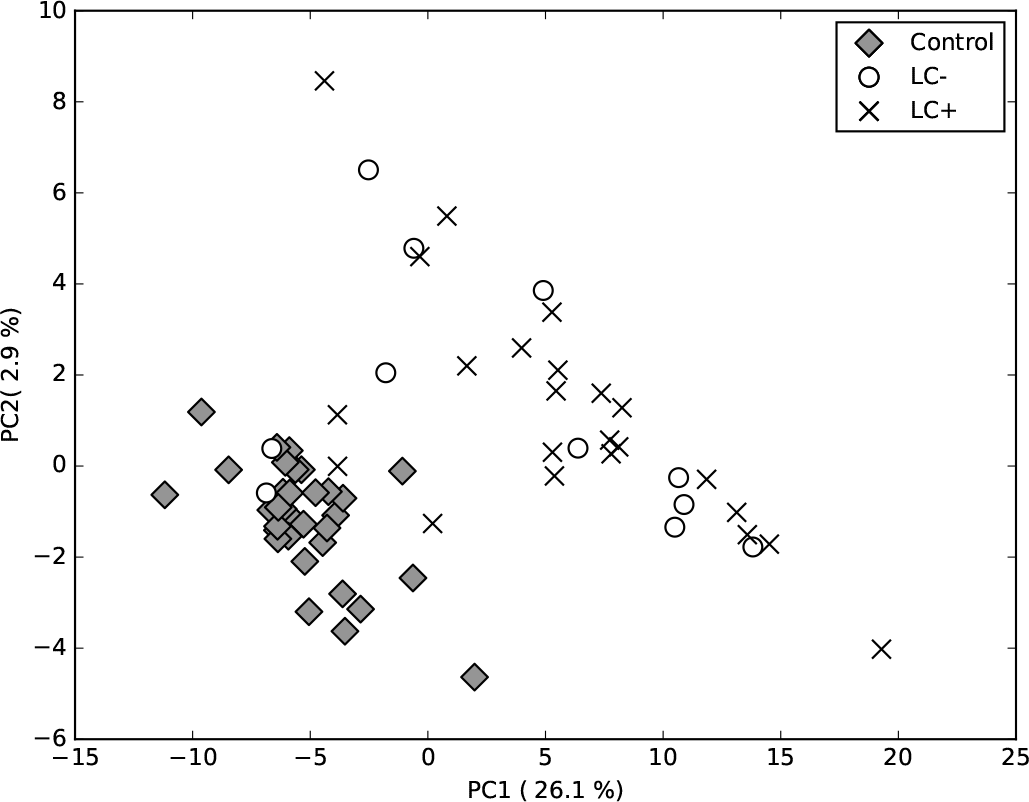
PCA score plots of FIE-MS data using all normalised *m*/*z* intensity values in negative ionisation mode, showing no clear separation between LC+ and LC− samples.

### 3.1 Analysis of small-cell lung cancer and non-small cell lung cancer

Determining the type of lung cancer that has developed in a patient is a key component of determining the correct treatment and management pathway. For lung cancer, two broad classes of classification exist: non-small-cell lung cancer (NSCLC) and small-cell lung cancer (SCLC). Patients with NSCLC are usually classified as one of three main subtypes: adenocarcinoma, squamous-cell carcinoma and large-cell carcinoma. Of these, adenocarcinoma is the most common and is characterised by overproduction of mucin. Squamous-cell carcinoma is the second most common form of lung cancer and typically occurs in the centre of the lungs. Large-cell carcinoma is less common and is characterised by cancerous cells that are large, with excess cytoplasm and large nuclei. The extent of NSCLCs is reported using the TNM format, which is important for prognosis and treatment planning. The TNM format ranges from Stage 0 to Stage IV, with the relevant stage determined through assessment of the primary tumour, involvement of regional lymph nodes, and the extent of distant metastasis against set criteria [19].

**Figure 4:**
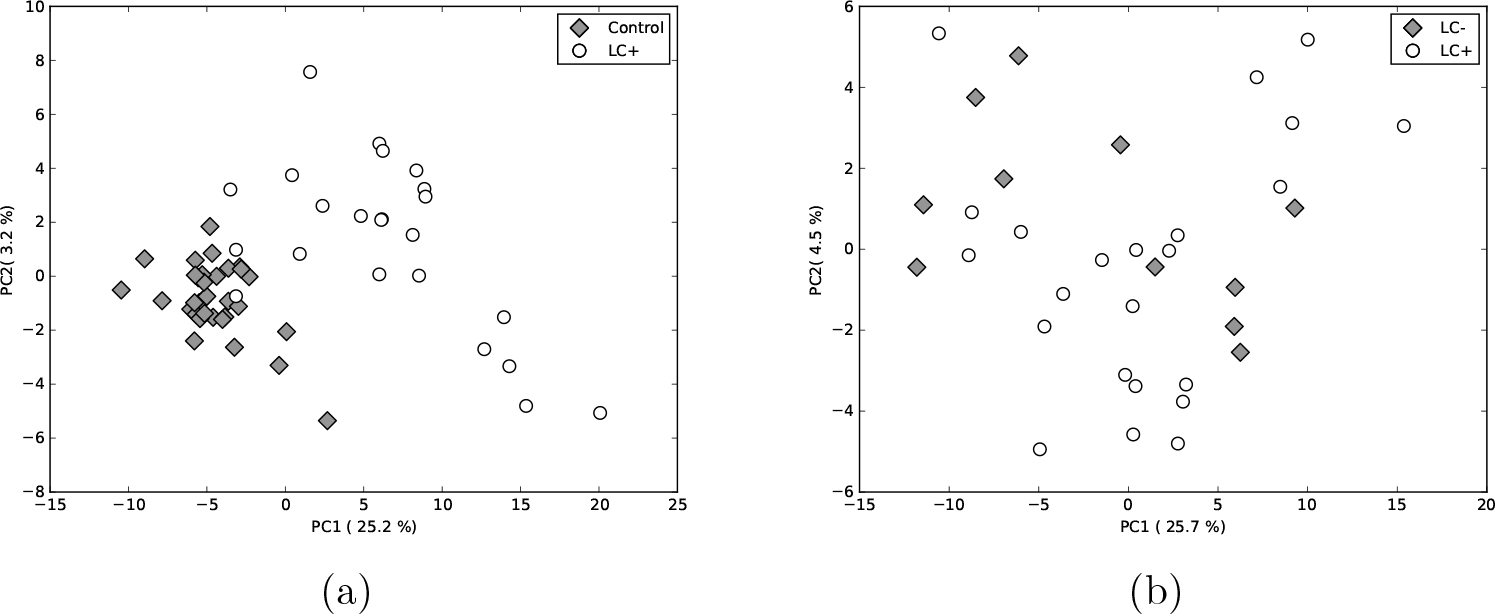
PCA score plots of the FIE-MS data with (a) only 970 selected *m*/*z* signals for LC+ and healthy controls, and (b) 125 selected *m*/*z* signal for LC+ and LC-. Using *m*/*z* features taken from *t*-tests clearly differentiates between relevant classes.

Small-cell lung cancers are less common than NSCLC, with approximately 10% of all lung cancers classified as SCLC. These lung cancers are characterised by their small cells, with minimal cytoplasm, and poorly-defined cell borders. Cancerous cells are usually rounded, oval and spindle-shaped. Typically, patients with SCLC present when the disease has metastasised from the lungs and symptoms frequently this, such as issues with bone marrow and the liver because of metastasis. Small-cell lung cancers are staged differently to NSCLC. Although the TNM format can be used, it does not predict survival and other outcomes well. Typically, SCLC is staged as either limited or extensive disease, with the latter equivalent to Stage IV of the TNM staging format for NSCLC [19].

**Figure 5:**
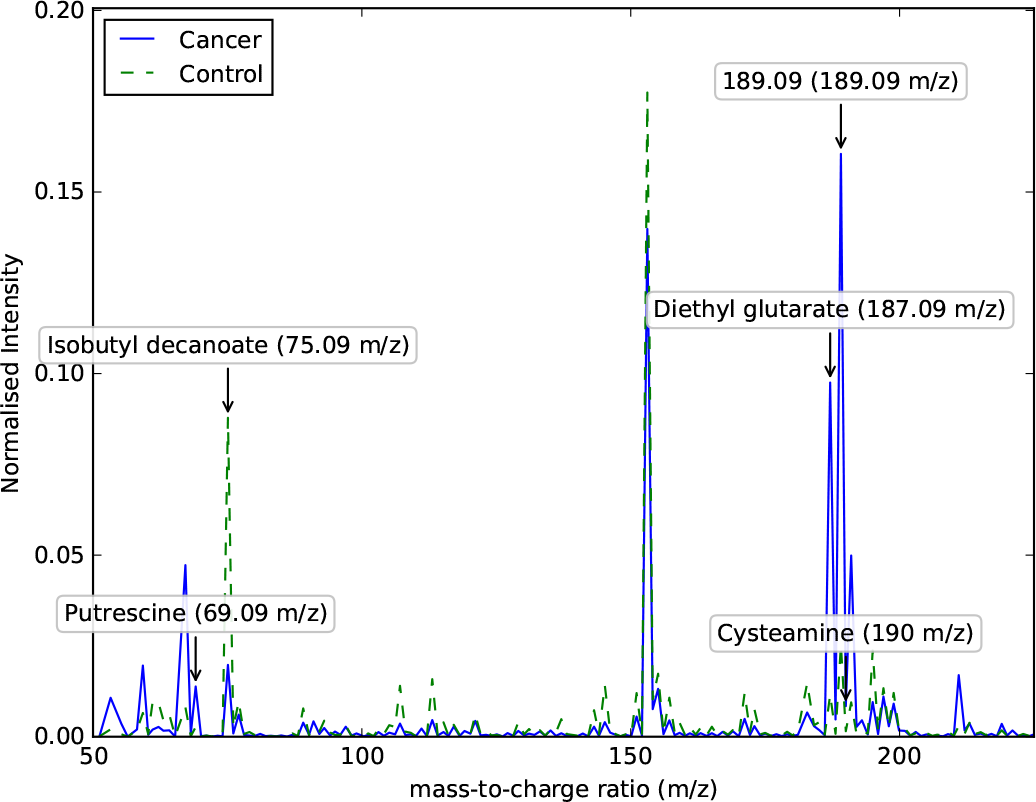
Mean FIE-MS spectra illustrating five key distinguishable metabolites between patients with lung cancer (LC) and healthy controls.

**Figure 6:**
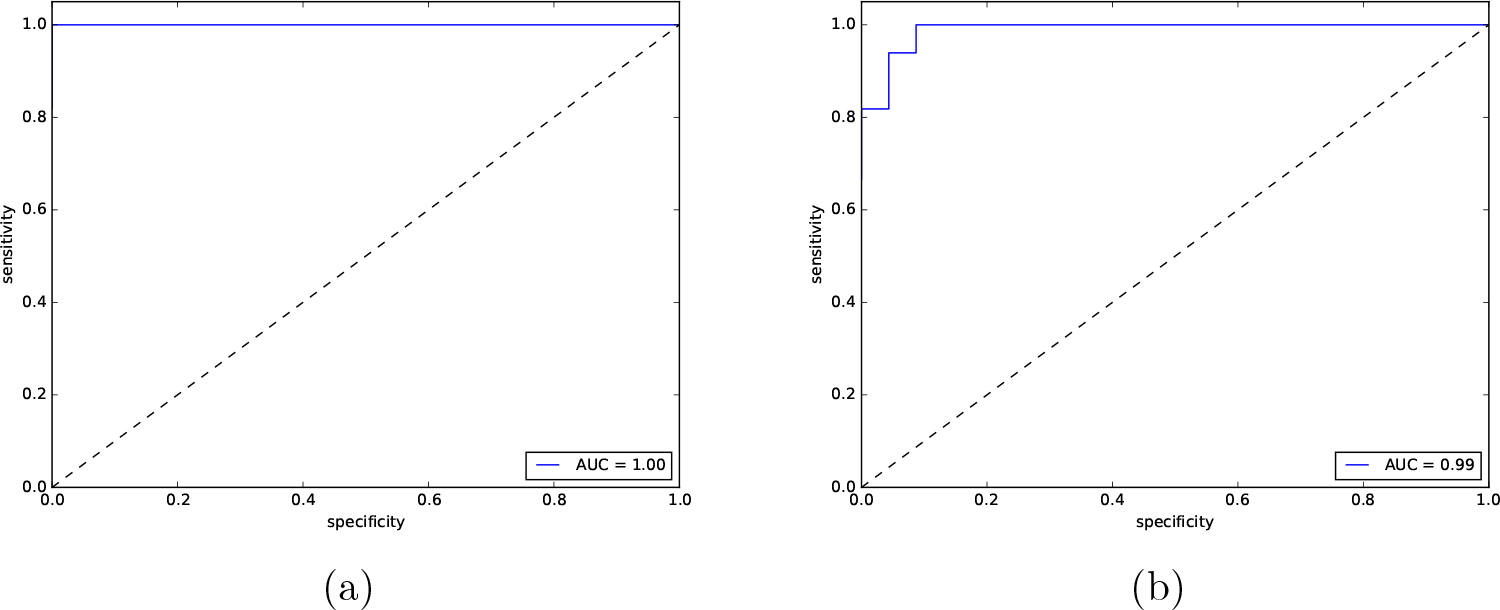
Receiver operating characteristic curves obtained from ANN models using leave one out cross-validation for classifying: (a) LC+ against LC−, and (b) LC against healthy controls.

A total of nine *m*/*z* values provided strong differentiation between SCLC and NSCLC with *p*-values less than 0.05 from Welch *t*-tests. Out of these 9 *m*/*z* values, 6 were identified and brought forward for further analysis as potential biomarkers of NSCLC and SCLC. Their relative values and *p*-values are shown in Table 3. Furthermore, ranges of each metabolite shown as box- and-whisker plots can be found in Figure 7. These both indicate that the 6 biomarker candidates show that the median levels of the 6 markers are higher in NSCLC samples to patients with SCLC.

**Table 3:** Identified metabolites that are significantly different between patients with Non-Small Cell Lung Cancer (NSCLC) and Small Cell Lung Cancer (SCLC) using Welch *t*-tests.

| Metabolite | $m/z$ | Normalised Intensity | | $p$-value |
| --- | --- | --- | --- | --- |
|  |  | NSCLC | SCLC |  |
| Phenylacetic acid | 137.09 | $-3.31 \pm 0.21$ | $-2.85 \pm 0.42$ | 0.001 |
| L-Fucose | 165.09 | $-3.19 \pm 0.12$ | $-2.86 \pm 0.30$ | 0.001 |
| Caprylic acid | 145.18 | $-2.81 \pm 0.44$ | $-2.12 \pm 0.31$ | 0.001 |
| Acetic acid | 61.09 | $-2.96 \pm 0.19$ | $-2.56 \pm 0.40$ | 0.002 |
| Propionic acid | 75.09 | $-1.91 \pm 0.18$ | $-1.56 \pm 0.34$ | 0.003 |
| Glycine | 76.09 | $-3.17 \pm 0.12$ | $-2.91 \pm 0.21$ | 0.004 |

PCA analysis (Figure 8a) showed good separation capabilities between NSCLC and SCLC samples, 6 metabolites were selected as input features to build a second ANN. Due to the small sample size, leave-one-out cross validation was performed to estimate the generalisation performance of the model. Our MLP model was able to distinguish between NSCLC and SCLC with a sensitivity of 80% and a specificity of 100% for predicting SCLC from cross-validation (see Table 2 and Figure 8b).

## 4. Biomarker analysis

Metabolic profiling recognised and provided identifications of 6 candidate metabolites that offered superb predictive values. Amongst the targeted metabolites are examples which have already been linked to lung cancer. The enzyme glycine decarboxylase (GLDC) is involved in the degradation of glycine which is coupled to the generation of methylgroups which can be used in (for example) purine biosynthesis. GLDC expression was increased in cells isolated from NSCLC tumours with concomitent decreases in glycine [20]. These authors showed that GLDC expression could serve as a biomarker, we now provide evidence that relative decreases in glycine is a feature of NSCLC in sputum. This biofluid represents a less invasive and potentially cost-effective means of clinically assessing patient LC status.

**Figure 7:**
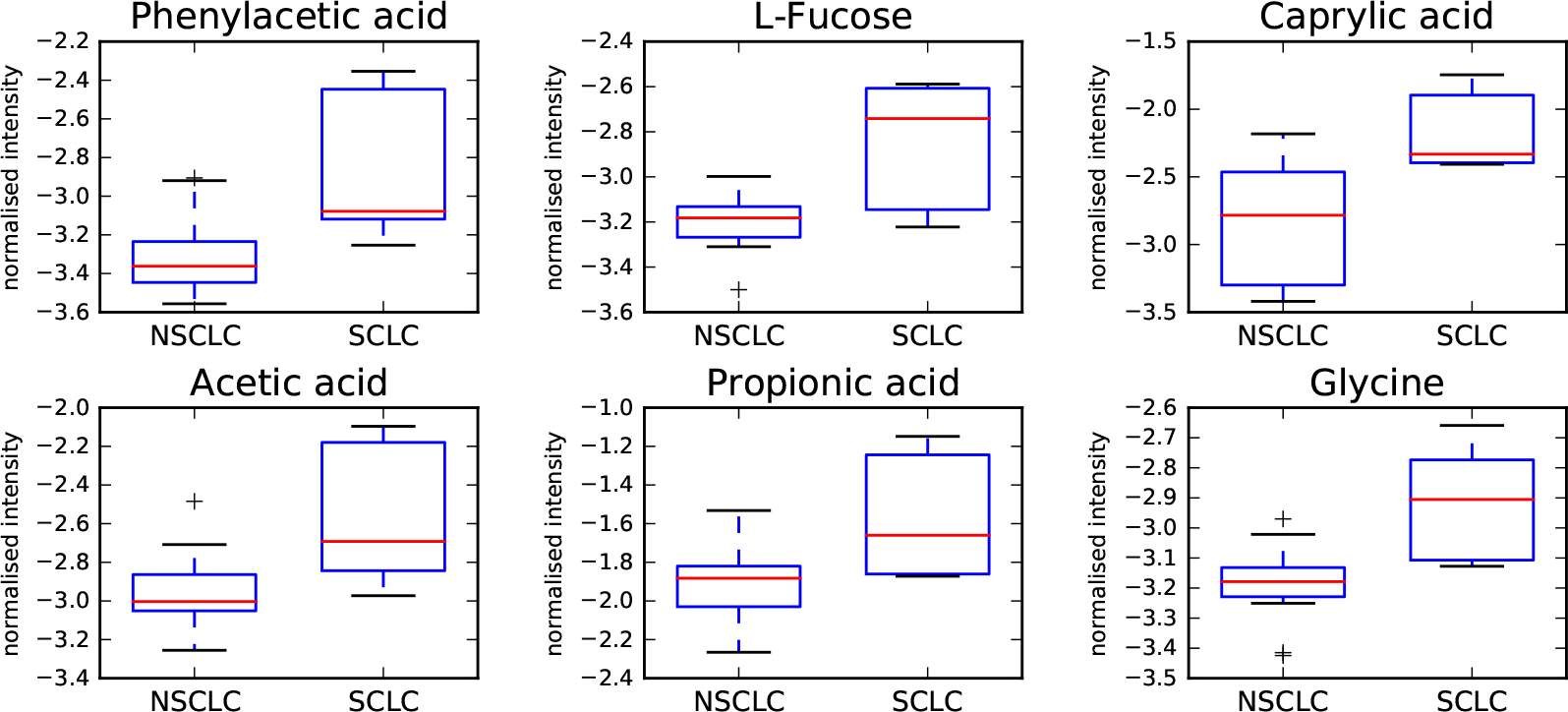
Box-and-whisker plots of the candidate metabolite biomarkers for discrimination between NSCLC and SCLC. The y axis represents the normalised intensity of each metabolite. Horizontal lines in the middle portion of the box illustrates the median, bottom and top boundaries of boxes represent the lower and upper quartile, whiskers depict the 5th and 95th percentiles and plus signs depicts the outliers.

Fucose (6-deoxy-L-galactose) is N- and O-linked to a range of glycolipids and glyocpeptides produced by mammalian cells. Increases in fucoslylated proteins, for example, a-fetoprotein are used in the diagnosis of hepatocellular carcinoma [21]. Fucosylation is dependent on the availability of the substrate guanosine 5’-diphospho-fucose (GDP-Fucose) and associated glycosyltransferases to transfer the fucose motif on to the protein / lipid. Fucose is a precursor to GDP-Fucose production [22] so that increases in fucoslyation could lead to a relative depletion in fucose as noted in our study. Increased fucosylation has been previously linked to NSCLC biopies [23] but ours is the first suggestion that decreases in fucose pools in sputum could be clinically suitable marker.

**Figure 8:**
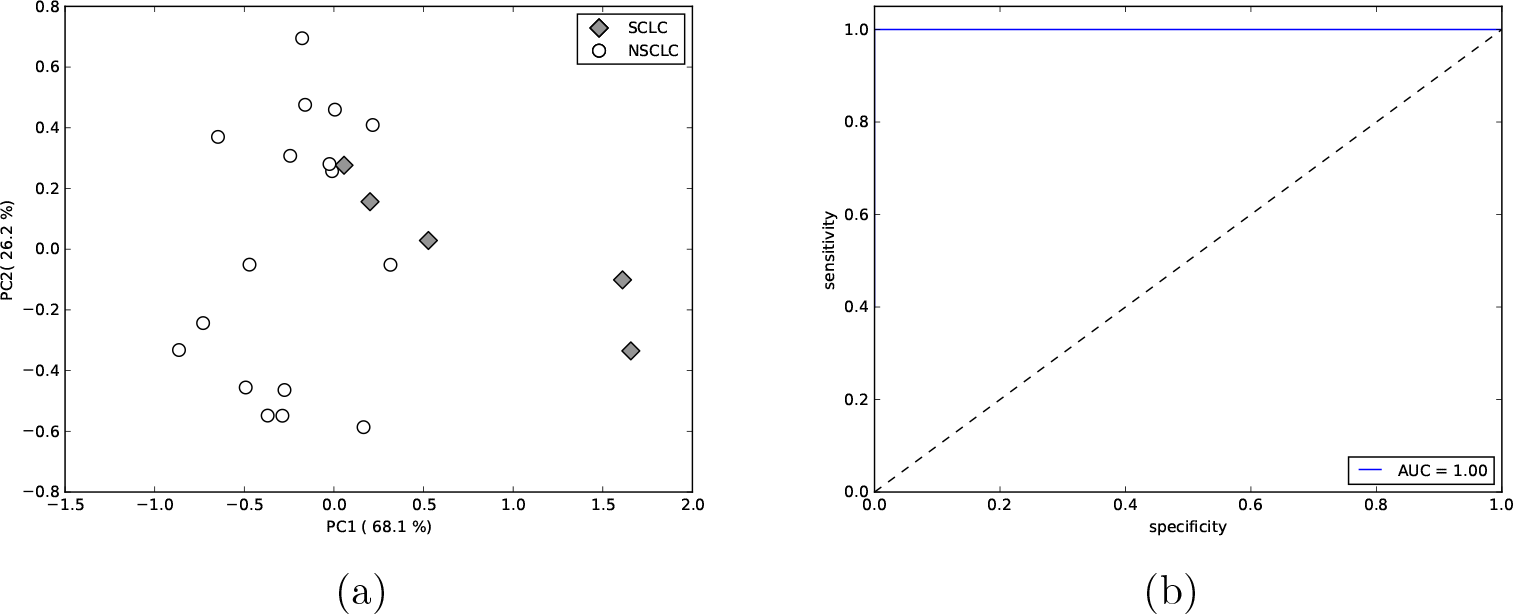
PCA score plot of the 6 identified metabolites for (a) NSCLC and SCLC, and (b) the ROC curve obtained from ANN LOOCV.

Other key metabolites were volatile short chain fatty s acetic (C2), propionic (C3) and caprylic [octanoic] (C8), These could be derived as a result of lipid peroxidation [24] but irrespective of their means of generation would provide further support for efforts that are attempting to sample breath as an non-invasive method for lung cancer detection [24, 25, 26].

A somewhat surprising observation was the detection of phenylacetic. This is classically associated with phenylketonuria (PKU); an inherited disorder of amino metabolism. PKU arises from a deficiency of the liver enzyme phenylalanine-4-hydroxylase which production of tyrosine from phenylalanine. If this enzyme is non-functional a range of alternative metabolites are produced, including phenylacetic [27]. However, beyond PKU, phenylacetate accumulates in patients with chronic kidney disease and during renal failure where it can inhibit nitric oxide generation [28] and macrophage intracellular killing of bacteria [29]. Further, disease-associated phenylacetate accumulation can contribute to an inflammatory responses [30]. To our knowledge, phenylacetate has not been associated with cancer; indeed quite the opposite it has a history of being tested for its anti-tumour properties [31]. However, it may be that NSCLC has particularly phenotypic / biochemical features which lead to altered phenylalanine metabolism leading to the accumulation of phenylacetate to contribute to inflammatory events and a reduced ability to deal with the lung microbiome. This is currently under investigation in our group.

## 5. Conclusions

A metabolomics approach based on FIE-MS coupled with univariate Welch *t*-test based feature selection and artificial neural networks provides an efficient methodology for metabolomic-based profiling of sputum to differentiate between non-small cell and small-cell lung cancer. This paper has identified 6 candidate metabolites markers, including L-fucose, phenylacetic, caprylic, acetic, propionic acid, and glycine, which were found to have good discriminatory abilities and low *p*-values. Excellent sensitivity and specificity was also shown using these markers through leave-one-out cross validation, which further indicates the promise of metabolomic analysis of sputum for non-invasive screening for LC. Further analysis involving a larger number of samples is required to determine both the precision and applicability of this approach in guiding the diagnosis and treatment of LC and respective forms.

## Acknowledgements

The work described in this paper was funded by two Aberystwyth University PhD scholarships to Keiron O’Shea and Simon Cameron. Keiron O’Shea would like to thank Nicholas Dimonaco and Manfred Beckmann of Aberystwyth University for their helpful support offered during this project.

